# A high-throughput delayed fluorescence method reveals underlying differences in the control of circadian rhythms in *Triticum aestivum* and *Brassica napus*

**DOI:** 10.1101/570572

**Authors:** Hannah Rees, Susan Duncan, Peter Gould, Rachel Wells, Mark Greenwood, Thomas Brabbs, Anthony Hall

## Abstract

**Background:** A robust circadian clock has been implicated in plant resilience, resource-use efficiency, competitive growth and yield. A huge number of physiological processes are under circadian control in plants including: responses to biotic and abiotic stresses; flowering time; plant metabolism; and mineral uptake. Understanding how the clock functions in crops such as *Triticum aestivum* (bread wheat) and *Brassica napus* (oilseed rape) therefore has great agricultural potential. Delayed fluorescence (DF) imaging has been shown to be applicable to a wide range of plant species and requires no genetic transformation. Although DF has been used to measure period length of both mutants and wild ecotypes of *Arabidopsis*, this assay has never been systematically optimised for crop plants. The physical size of both *B. napus* and *T. aestivum* led us to develop a representative sampling strategy which enables high-throughput imaging of these crops.

**Results:** In this study, we describe the plant-specific optimisation of DF imaging to obtain reliable circadian phenotypes with the robustness and reproducibility to detect diverging periods between cultivars of the same species. We find that the age of plant material, light regime and temperature conditions all significantly effect DF rhythms and describe the optimal conditions for measuring robust rhythms in each species. We also show that sections of leaf can be used to obtain period estimates with improved throughput for larger sample size experiments.

**Conclusions:** We present an optimized protocol for high-throughput phenotyping of circadian period specific to two economically valuable crop plants. Application of this method revealed significant differences between the periods of several widely grown elite cultivars. This method also identified intriguing differential responses of circadian rhythms in *T. aestivum* compared to *B. napus*; specifically the dramatic change to rhythm robustness when plants were imaged under constant light versus constant darkness. This points towards diverging networks underling circadian control in these two species.

## Background

A circadian clock is an endogenous oscillator entrained by external temporal cues. Circadian control of gene expression is a ubiquitous feature which appears to have arisen independently in bacteria, fungi, plants and animals(1). Since the discovery of the first *Arabidopsis* circadian mutant in 1995(2), the significance of the circadian clock in plants has become increasingly evident. Approximately 30% of genes in *Arabidopsis* are predicted to be under circadian control, regulating photosynthetic, metabolic and developmental pathways(3,4). Moreover, a selective advantage resulting from a clock which is matched to the exogenous day-length has been demonstrated in mammals, insects, bacteria and plants(5-9).

The most recent model for the molecular control of the *Arabidopsis* clock is comprised of a series of interlocking negative transcriptional feedback loops regulated by key activators which control the oscillation of clock gene expression (10). To ascertain the underlying nature of circadian rhythms, a clock-controlled output representing the pace of the clock must be measured in constant (free-running) conditions. Previously this research has been conducted by studying leaf movement rhythms or by following luciferase gene expression under the control of a circadian regulated promoter(11-13). Delayed fluorescence (DF) imaging provides an alternative to these methods that does not require plant transformation. It has previously been shown to work in a variety of plants for which leaf movement assays are not feasible (14,15). Delayed fluorescence occurs when excited electrons in photosystem II (PSII) undergo spin-conversion to a triplet excited state before charge recombination allows them to return to their ground state releasing light energy(16). Measurements of DF have been correlated with the photosynthetic state of PSII(17) and the amount of DF production is regulated by the circadian clock. DF can be measured with a low-light imaging system identical to that used for luciferase imaging and output rhythms have been shown to oscillate with a comparable period to those estimated from luciferase reporter experiments(14). The output from a DF experiment is a waveform which has parameters that can be mathematically defined and therefore quantified. These parameters include ‘period’ (the time taken to complete one cycle), ‘phase’ (the time of day at which this peaks) and ‘amplitude’ (the distance between the peak and the baseline of the oscillation). Important to circadian dynamics is also the idea of ‘rhythm robustness’ i.e. whether these parameters change overtime. In this paper, rhythm robustness was assessed by: the percentage of samples classified as rhythmic; the relative amplitude error (RAE); the period CV and the average period error threshold (all defined in Supplementary Materials). Together, these parameters allow the effects of different imaging conditions to be quantified.

As DF measurement is correlated with the oscillations in photosynthetic status of PSII, leaf material is the logical choice for a representative sample. Rhythms have been shown to persist in excised leaves in several species(18-22). However, previous research has demonstrated that independent clocks run at different periods throughout the plant under constant conditions, coordinated by a degree of intercellular coupling(23-27). The extent to which the clock is affected by dissecting leaf material into small segments is investigated in this paper.

Alongside these spatial differences, the clock has also been shown to be temporally dynamic and is affected by the life history of the plant. Both the systemic age of the plant and the ‘emergence age’ of the individual leaves on a plant have been reported to effect the clock in *Arabidopsis*, with increasing age associated with period reduction(28). Conversely, the timing of leaf senescence has also been shown to be directly regulated by core circadian genes(29).

In addition to this endogenous entrainment, the clock is also responsive to external stimuli; the most well characterized of which are light and temperature cues. Increasing light intensity causes a shortening of period in free-running conditions(30-32) and these rhythms rapidly dampen in amplitude under continuous darkness(33). Circadian systems are relatively buffered against temperature changes compared to other biochemical reactions but are not completely independent of it(34). Period shortening of 1.8-4.2h have been reported following temperature increases from 17°C to 27°C determined by both leaf-movement assays and luciferase reporters under circadian regulation in *Arabidopsis(31,35,36).* Seedlings grown at 17°C also have rhythms with lower period variability and RAE values than plants grown at 27°C(35,37). The extent to which rhythms are temperature compensated is described using the inverse of the temperature coefficient Q10; the change in the rate of a process over a temperature change of 10°C(38).

Here we present an optimized protocol for high-throughput phenotyping of circadian period using two crop plant models; *Triticum aestivum* (bread wheat) and *Brassica napus* (oilseed rape). Both *T. aestivum* and *B. napus* are polyploid organisms that have been influenced by human domestication, genome duplication events and geographical speciation as the use of these crops became globalized. The specific and combined effects of these factors on the control of the clock is yet to be investigated.

Within this paper we show that both the age of the plant and the developmental age of leaves have significant effects on period with older material displaying shorter rhythms. *Brassica* and wheat seedlings exhibited a shortening of period of approximately half an hour to almost three hours per week after sowing. Period differences of up to 3.14h were observed between leaves of plants of a single systemic age. To make our method high-throughput whilst still providing reliable rhythms, tissues were segmented into various sizes and compared to whole leaf samples. We identify regions of the plant leaves which are the most robustly rhythmic and give the most consistent period estimates.

Both the light regime and temperature conditions also had large effects on period estimation. Constant light (L:L) produced robust rhythms in *B. napus* and constant darkness (D:D) produced more robust rhythms in *T. aestivum.* Warmer temperatures accelerated DF rhythms in both *Brassicas* and wheat, indicating a curtailed temperature compensation response. Intriguingly, we also demonstrate that rhythm robustness is positively correlated with increasing temperature in wheat; opposite to what has previously been observed in *Arabidopsis* (29).

Finally, we applied our optimized, high-throughput DF method to investigate differences between elite cultivars in both *B. napus* and *T. aestivum* and demonstrate it to be a useful tool for assaying circadian rhythms in these crop species.

## Results

### Circadian variability due to leaf development and age of plant

We tested the effect of both plant and leaf aging on period estimates from *Brassica* and wheat seedlings. Previous studies in *Arabidopsis* have reported that the pace of the clock increases as the plant ages and that earlier emerged leaves have a shorter period than those which emerge later within the same individual(28). Our results mirror these findings for young wheat and *Brassica* plants, however this association was lost for older material (Figure 1A). For wheat we calculated period estimates from the second leaf of plants at 18, 25, 32 and 39 days after sowing and show that between 18-32 days period decreases linearly at a rate of approximately half an hour per week while maintaining a near constant relative amplitude error (RAE) (Figure 1A-C). However, in leaves from 39 day old plants there was an increase in both average period and relative amplitude error, potentially due to metabolic changes as a leaf changes from a source to a sink tissue or due to the onset of senescence in these samples. A one-way analysis of variance yielded a significant effect of wheat age on both period and RAE (F(3,90)=12.13, p<0.001) and (F(3,90)=7.018, p<0.001) respectively. Based on our investigation, we recommend using plants between 25 to 32 days after sowing. At 25 days 100% of samples were classified as rhythmic and period CV was 1.52h. 32 day old samples were also robust, having the lowest RAE (0.15) and period error (0.43) averages (see Supplementary Materials S1.)

**Figure 1.**
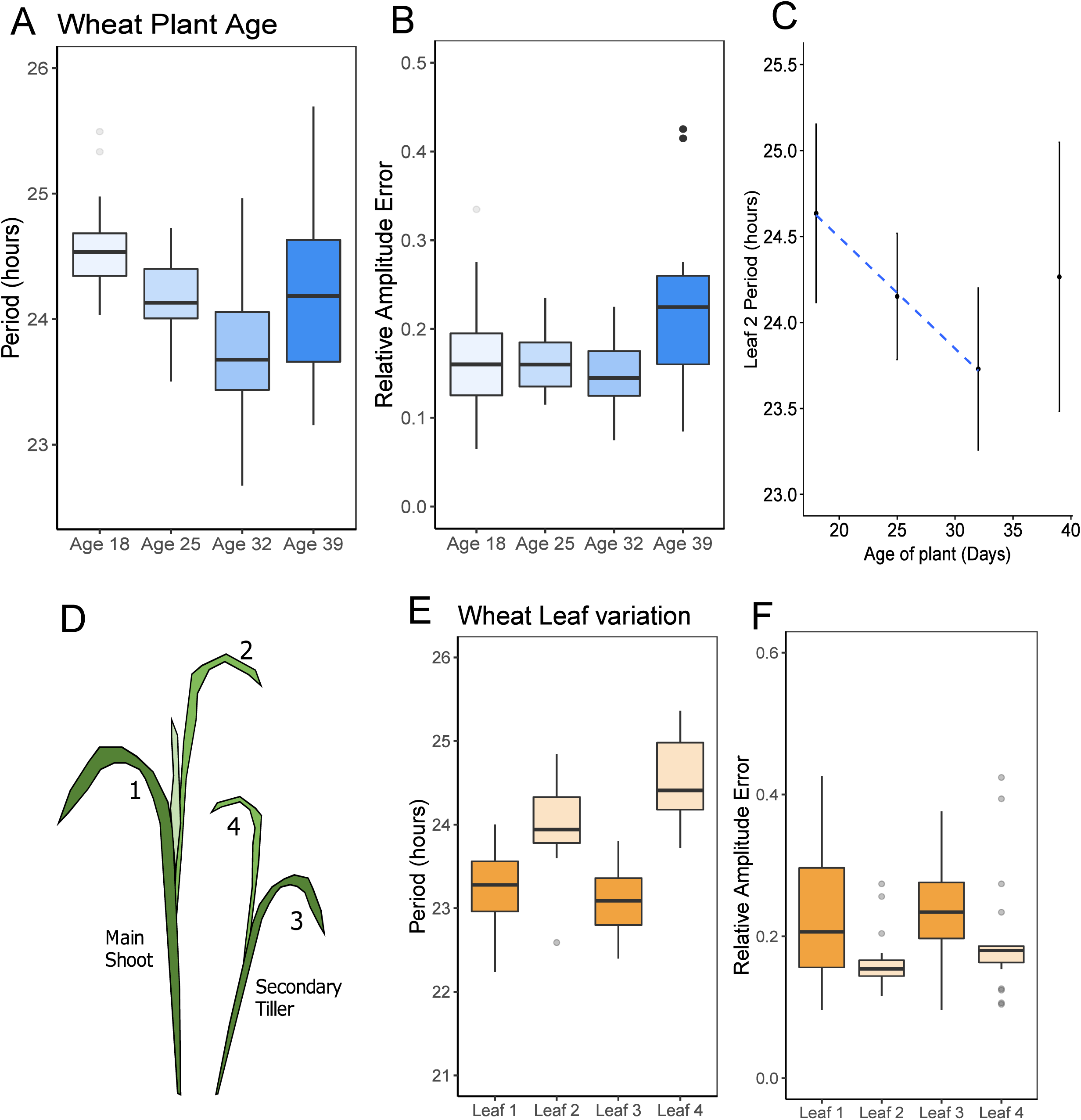
DF rhythms in wheat change with the age of the plant and between leaves on the same plant. The wheat plant age experiment (A-B) **used ‘leaf 2’ from plants grown for 18**, 25, 32 or 39 days. Blue boxplots show differences in period (A) and RAE (B) at each plant age. Figure C shows the mean periods from which the increase in rate was approximated (error bars are standard deviations and the blue dashed line is a linear trend for first three ages). The wheat leaf variation experiment used 4 leaves sampled from 25 day old plants following the leaf numbering system described in C. Orange boxplots show differences in period (D) and RAE (E) at each leaf age. Colour scales reflect an ageing gradient with lighter colours representing younger material. Data represents results from two imaging cabinets run in parallel as technical replicates and normalised for the between-cabinet effects. Period estimates were calculated using FFT-NLLS (BAMP de-trended data, 24-120h cut-off). N values reflect the number of samples for which period was estimated out of the total number of individuals sampled. Age 18 (N=26/26), age 25 (N=24/24), age 32 (N=25/26), age 39 (N=19/23). Leaf 1 (N=22/22), leaf 2 (N=22/22), leaf 3 (N=22/22), leaf 4 (N=21/22). Significance codes: \*\*\**p*<0.001, \*\**p*<0.01 \**p*<0.05.

In a separate experiment, we analyzed 4 leaves from 25 day old wheat plants as is shown in Figure 1D, where leaves 1 and 3 were the oldest leaves and leaves 2 and 4 the second oldest leaves from the main and secondary tiller, respectively. There was a statistically significant difference between the mean periods at each leaf age (one-way ANOVA (F(3,83)=7.434, p<0.001). Within each tiller pair, the older leaf had a shorter period than the younger leaf and had higher RAE averages. The mean period for leaf 4 (24.50h) was found to be significantly longer than both leaf 1 (23.15h) and leaf 3 (23.32h) (Tukey HSD). We recommend using leaf 2 as it had the best overall circadian robustness with regards to the % samples returned (100%), RAE (0.18) and period error (0.50) (Supplementary Materials S2).

For *Brassica* seedlings, plants were grown to 4 different ages: 20, 25, 30 and 35 days after sowing and leaf 1, 3 and 5 were sampled in the same experiment with leaf 1 being the earliest emerged leaf and leaf 5 the most recently emerged leaf (Figure 2). We conducted a nested ANOVA to test the effects of both plant age and within-plant leaf-age on period. We found that variation in plant-age had a significant effect on period, with increasing age causing a shortening of period (F(3, 53)=8.48, p<0.001). The nested effect of leaf age within each plant age-group was also found to be significant (F(8, 53)=5.45, p<0.001). The largest difference between leaves in each age-group was seen for 20 day old plants where a difference of 3.14h was observed between leaf 1 and 5 (p<0.001, Tukey HSD) (Figure 2A). Brassica plant-age was also found to have a significant effect on RAE averages with younger plants having a lower mean RAE (F(3,53)=5.953, p<0.01) (Figure 2B). Supplementary Materials S3 shows robustness statistics for all plant ages and leaf-ages tested. We recommend using leaf 1 from 20 day old plants as they had the lowest RAE (0.15) and period error threshold (0.47).

**Figure 2.**
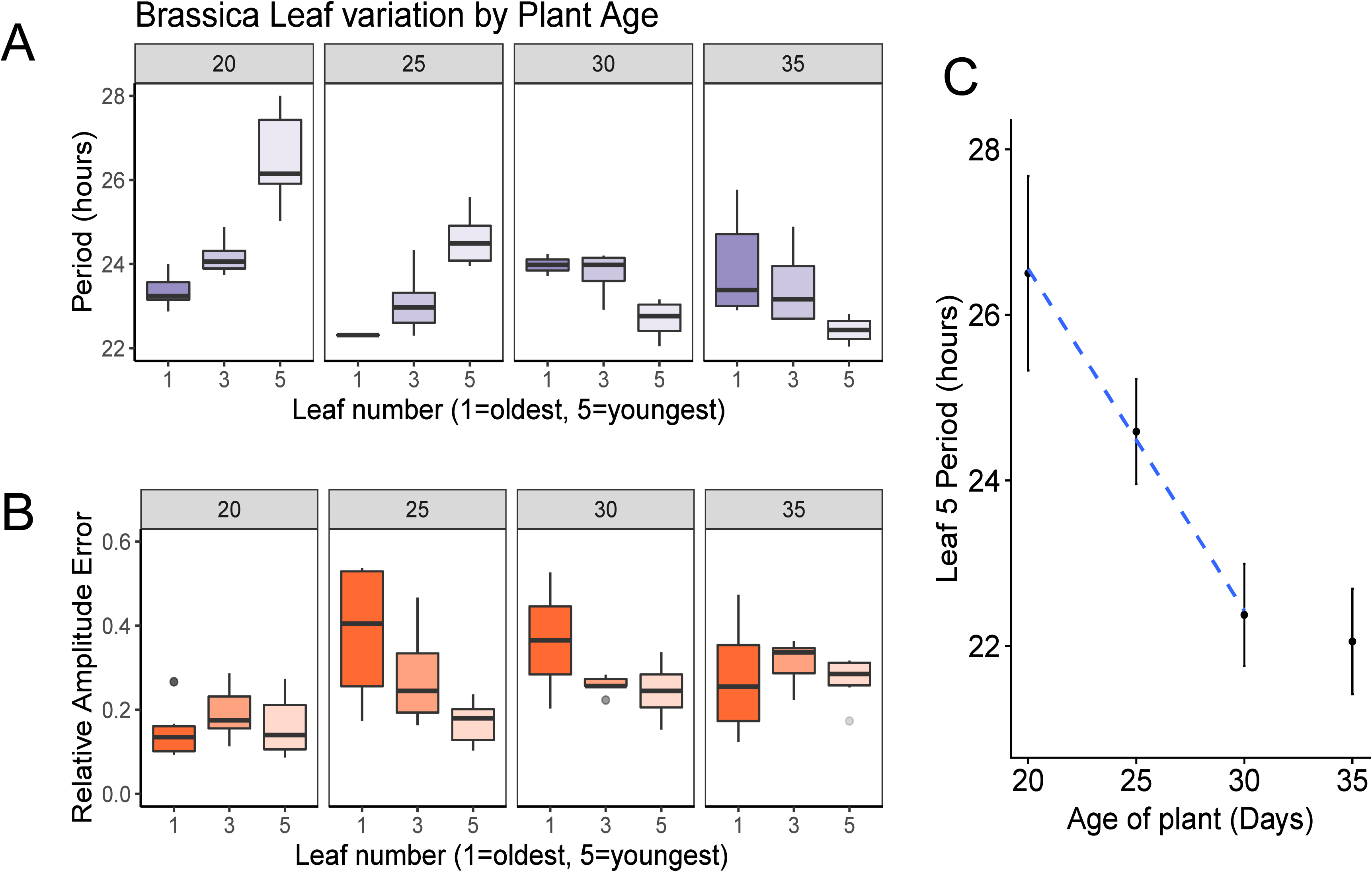
DF rhythms in *Brassica* change with the age of the plant and between different leaf ages. *Brassica* seedlings were grown to 4 different ages: 20, 25, 30 and 35 days after sowing Leaves 1, 3 and 5 were sampled from each plant in the experiment with leaf 1 being the earliest emerged leaf and leaf 5 the most recently emerged leaf. Boxplots show differences in period (A) and RAE (B) for each leaf age within each plant age. Colour scales reflect an ageing gradient with lighter colours representing younger material. Periods and RAE estimates were calculated using FFT NLLS (BAMP dtr, 24-120h cut-off). Data represents results from two imaging cabinets run in parallel as technical replicates and normalised for the between-cabinet effects. Age 20: leaf 1 (N=6/6), leaf 3 (N=6/6), leaf 5 (N=6/6). Age 25: leaf 1 (N=4/6), leaf 3 (N=6/6), leaf 5 (N=6/6). Age 30: leaf 1 (N=2/6), leaf 3 (N=5/6), leaf 5 (N=6/6). Age 35: leaf 1 (N=6/6), leaf 3 (N=6/6), leaf 5 (N=6/6). Significance codes: \*\*\**p*<0.001, \*\**p*<0.01 \**p*<0.05, all significance markers are relative to leaf 1 at each age.

To approximate the period shortening due to plant aging in *Brassica* we followed the changes in average period in leaf 5 across plant ages from 20 days old to 30 days old as is shown in Figure 2C. Our analysis revealed that period shortened by approximately 3 hours per week from a mean of 26.50h(SD 1.17) to 22.38h (SD 0.62).

### Finding an optimal size of leaf sample

We needed to identify representative leaf sections which allowed a sufficient number of samples to be analyzed on one plate without compromising the robustness of rhythms for period estimation. For wheat, we selected leaf 2 from 25 day old plants and analyzed the periods and circadian robustness given by whole leaves compared to leaves cut into 10cm sections and leaves cut into 4cm sections as shown in Figure 3A. By taking 4cm samples from 2 regions on the same leaf (5 or 15cm down from the tip) we could investigate changes in period across the length of the leaf. For *Brassica* seedlings we selected leaf 1 from 21 day old plants and then kept them whole, took 3cm square samples from the centre or quartered them (Figure 3B). This helped inform whether any changes in circadian characteristics were a result of size reduction or from sub-sectioning regions of the leaf. Our data showed that period and RAE averages were not significantly affected by cutting samples in wheat (Figure 3C and 3D), (F(3,75)= 2.066, p>0.1, one-way ANOVA). However, cutting *Brassica* leaves did significantly affect period estimates; quartered segments had a slightly longer period than whole samples (F(2,113)=5.46, p<0.01, one-way ANOVA, Tukey HSD (Whole-Quarter p<0.01) but RAE means were similar (Figure 3E and 3F) (Supplementary Materials S4). From this data we recommend using 10cm segments for wheat and 3cm square sections for *Brassica* imaging as these gave similar results to whole leaves and increased throughput by 44% for *Brassica* and 100% for wheat.

**Figure 3.**
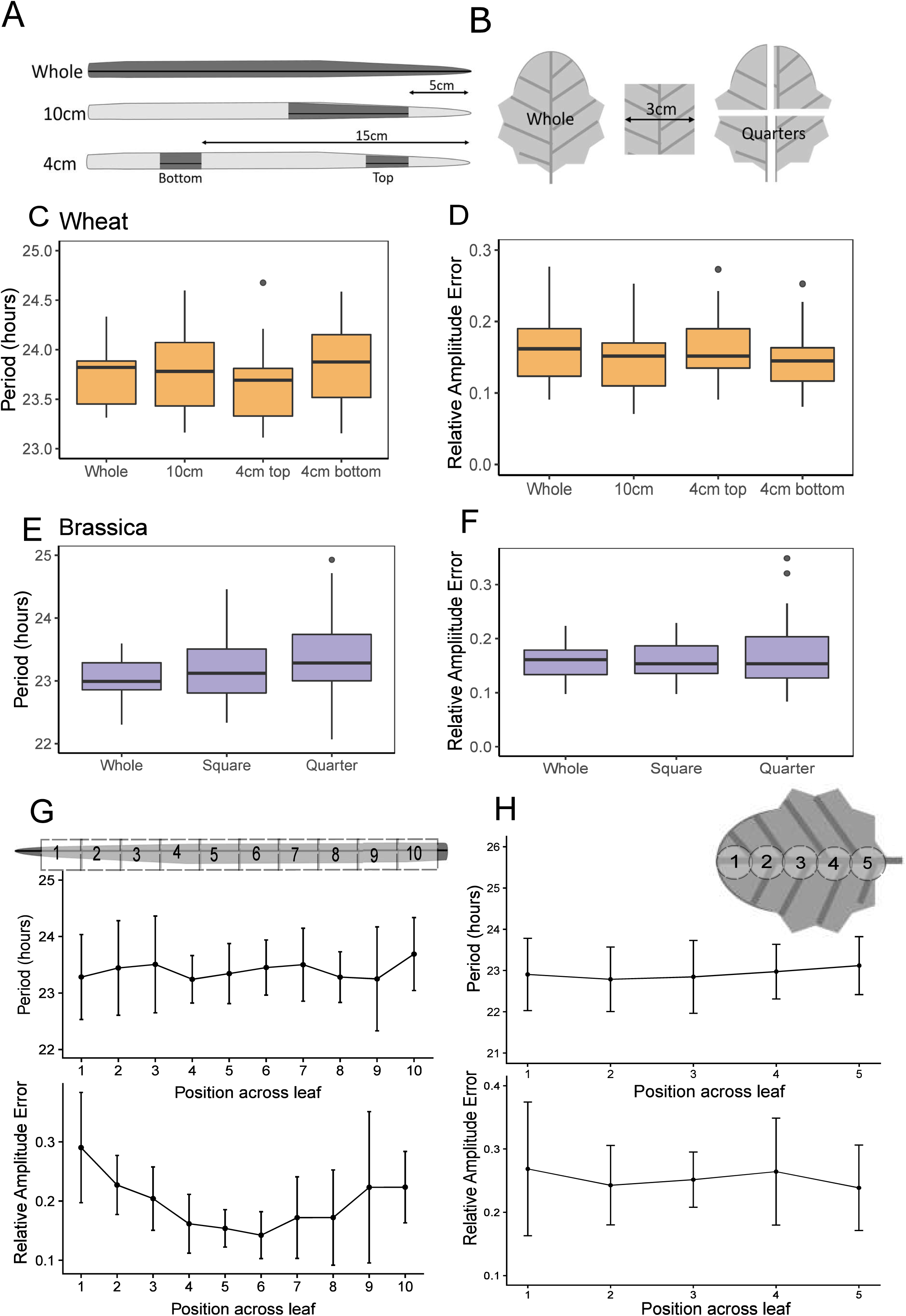
Cut sections of leaf material can be used to accurately make period estimates. The second leaf from 25 day old wheat seedlings was either left whole or sectioned into 10 cm or 4cm fragments cut either 5cm or 15cm from the tip (shown as dark grey sections in A). The first leaf from 21 day old *Brassica* plants was either left whole, sectioned into a 3cm square or quartered (B). Orange boxplots show differences in period (C) and RAE (D) for wheat sections. Purple boxplots show differences in period (E) and RAE (F) for brassica sections. Period and RAE were estimated using FFT NLLS, BAMP dtr, 24-120h cut-off. Whole leaves were digitally sectioned along the axis of the leaf post image-acquisition. Wheat period and RAE means for each section are shown in G. *Brassica* period and RAE means are plotted corresponding to the sectioning shown in H. Error bars show standard deviation. Data represents results from two experiments normalised for the between-experiment effects. Wheat: Whole (N=15/15), 10cm (N=23/23), 4cm top (N=20/21), 4cm bottom (N=22/22). *Brassica:* Whole (N=20/20), Square (N=21/21), Quarter (N=75/76). Significance codes: \*\**p*<0.01.

We next wanted to investigate whether period estimates changed across the axis of the leaf. We selected only the whole leaf images and digitally sectioned them into 10 or 5 regions of interest for wheat and *Brassica* leaves respectively (Figure 3G and 3H). Using this approach we observed an average within-leaf variance of 0.45h in wheat and 0.42h in *Brassica* leaves. This variation was larger than the leaf-to-leaf variation determined for wheat (0.04h) and *Brassica* (0.3h) leaves. The mean period and RAE for each section across these leaves was calculated and plotted (Figure 3G and 3H). No significant difference was observed between the period of wheat or *Brassica* segments; however the RAE was significantly different across wheat leaves (F(9,139)=6.077, p<0.001, One-way ANOVA). The middle segments (4, 5, 6, 7 and 8) had significantly lower RAE averages compared to the tip (segment 1) suggesting that this middle region may give the most robust DF rhythms.

### Constant free-running conditions: Dark versus Light

Two light regimes were tested which allowed free-running DF rhythms to be recorded. We entrained plants for 4 days in 12:12 light:dark (L:D) cycles at 22°C before sampling and imaging every hour under constant conditions, as described in Figure 4A. In both D:D and L:L conditions the exposure time was kept at 1 minute. Figure 4B shows the striking differences in period estimate accuracy obtained from wheat and *Brassica* under the two light regimes. For wheat, periods from leaves under a D:D regime had much lower variance than those under the L:L regime (D:D mean =23.29h, SD=0.53; L:L mean =23.54h, SD=3.19). For *Brassica* the opposite was observed; rhythms were more accurate under L:L (D:D mean=24.89h, SD=2.91; L:L mean=22.92h, SD=0.31). A shortening of period was observed for both *Brassica* and wheat under L:L compared to D:D based on median values, however the increased variance observed within wheat-L:L and *Brassica-*D:D resulted in these differences having low significance (wheat t(23.16)=-0.38, p>0.5; *Brassica* t(10.14)=2.24, p=0.048 Welch’s t-test).

**Figure 4.**
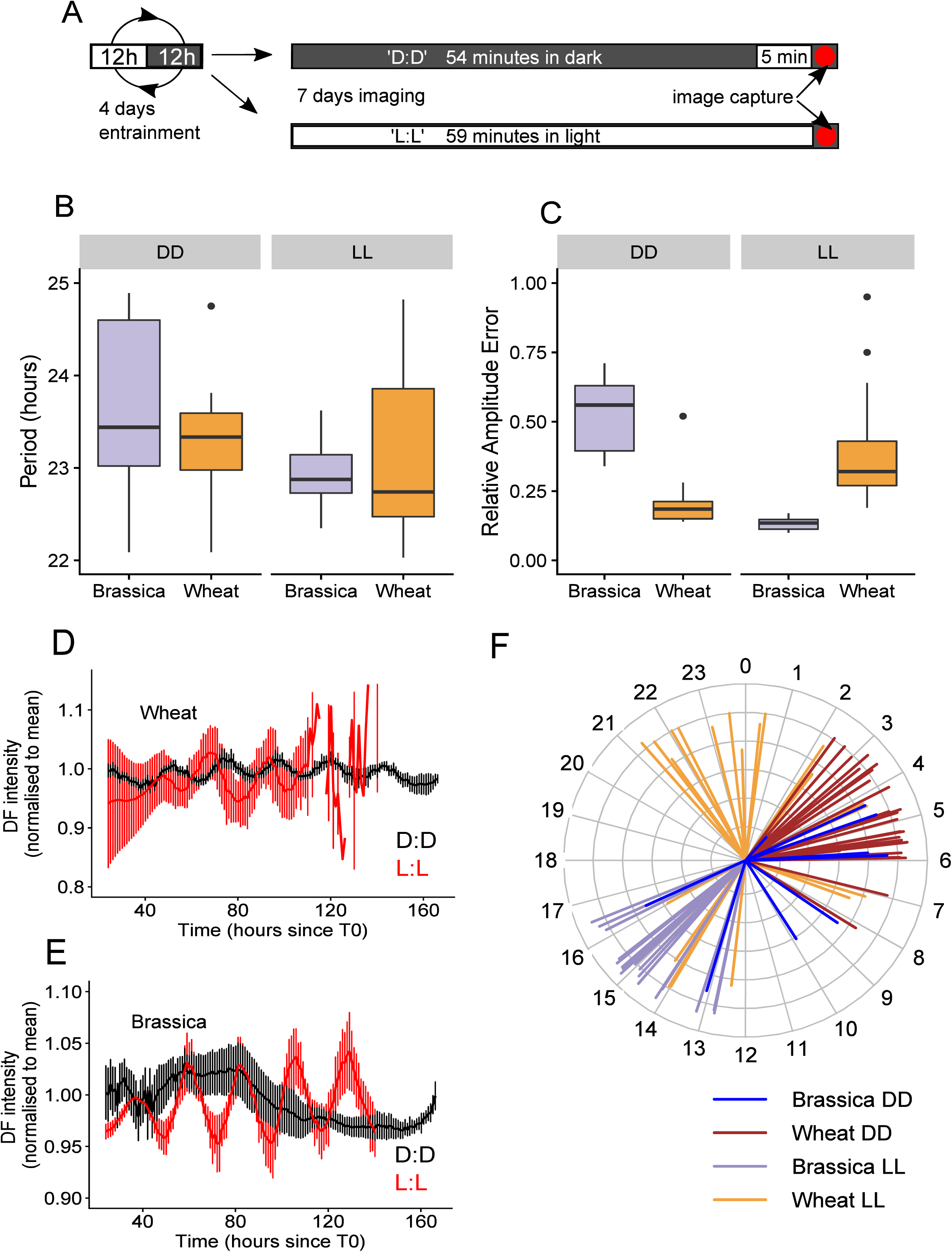
Effect of either L:L or D:D free-running light conditions on DF rhythms. Wheat and *Brassica* seedlings were entrained for 4 days in L:D at 22°C before sections were cut, plated and imaged. A D:D free run consisted of a loop of 54 minutes of darkness followed by 5 minutes of light exposure and then image capture. A L:L free-run consisted of 59 minutes of light exposure before image capture (A). Boxplots of period (B) and RAE (C) are shown for *Brassica* and wheat in D:D and L:L conditions where *Brassica* data is displayed in purple and wheat data in orange. Period and RAE were estimated using FFT NLLS, BAMP dtr, 24- 120h cut-off. Examples of oscillation traces are shown in D:D (black lines) and L:L (red lines) for wheat (D) and *Brassica* (E). Thick lines represent the mean trace of 6 mean-normalized individuals with error bars representing standard deviations. Estimated individual circadian phases are shown in the clock plot in (F) where the length of the line reflects the inverse circadian phase error (longer lines imply more confidence in the phase prediction). All phase estimates are relative to 0 where 0 represents entrainment dawn and 12 represents dusk. D:D *Brassica* (N=12/18); D:D Wheat (N=24/24); L:L *Brassica* (N=18/18); L:L Wheat (N=23/24). Data is consistent with additional preliminary experiments which can be seen in Supplementary S11 and S12. Significance codes: \*\*\**p*<0.001.

RAE ratios reflected the accuracy seen in period estimation between the regimes (Figure 4C). RAE averages were smaller in D:D for wheat (D:D mean=0.20, SD=0.08; L:L mean=0.38, SD=0.18) and in L:Lfor *Brassica* (D:D mean=0.53, SD=0.13; L:L mean=0.13, SD=0.02). RAE differences were significant between regimes for both species (Wheat t(29.23)=-4.38, p<0.001; *Brassica* t(10.39)=9.92, p<0.001 Welch’s t-test). Figure 4D and E show mean oscillation traces which demonstrate how DF rhythms were sustained in wheat and *Brassica* under the different light conditions. Interestingly, DF rhythms also had a dawn-phased peak in wheat and a dusk-phased peak in *Brassica* which became shifted as different light conditions were applied (Figure 4F).

Wheat samples under the D:D conditions returned 100% of samples from period estimation (L:L=95.83%), an average RAE ratio of 0.20 (L:L=0.38), a period CV of 2.25% (L:L=13.53%) and a period error threshold of 0.55 (L:L=1.17). For *Brassica*, rhythms under the L:L regime returned 100% from period estimation (D:D=66.67%), a RAE average of 0.13 (D:D=0.53), a period CV of 1.34% (D:D= 11.80) and a period error threshold of 0.41 (D:D=1.62). See Supplementary Materials S5. We would therefore recommend running wheat DF experiments under D:D conditions and *Brassica* DF experiments under L:L.

### Finding an optimum free-running temperature

To investigate the effect of temperature on period and rhythm robustness we tested *Brassica* and wheat seedlings at a range of constant temperatures. We used the optimal conditions from the variables so far tested and entrained each batch of plants at the imaging temperature for four days prior to imaging (see Methods). Both *Brassica* and wheat experienced an acceleration of the clock at higher temperatures, with the rate increasing most dramatically at lower temperatures (Figure 5A). Periods decreased from 26.40h (SD=3.60) at 17°C to 22.48h (SD=0.31) at 32°C in wheat. Periods decreased from 26.28h (SD=0.72) at 12°C to 23.16h (SD=0.52) at 22°C in *Brassica.* The temperature coefficient Q10 was calculated as an average across all temperatures (Supplementary Materials S7). Q10 was found to be 1.12 for wheat and 1.14 for *Brassica* indicating a degree of thermal compensation, but to a lesser extent than has been previously reported in *Arabidopsis* (36). We next looked at which temperatures gave the best rhythmicity in each crop. Rhythms were most robust in wheat grown at 27°C: 100% of period estimates were returned, the average RAE ratio was 0.15, period CV was 2.48% and period error threshold was 0.48 (Supplementary S6). There was a clear negative trend in period CV as the temperature increased in wheat from 13.63% at 17°C to 1.38% at 32°C. Mean RAE at each temperature can be seen in Figure 5B.

**Figure 5.**
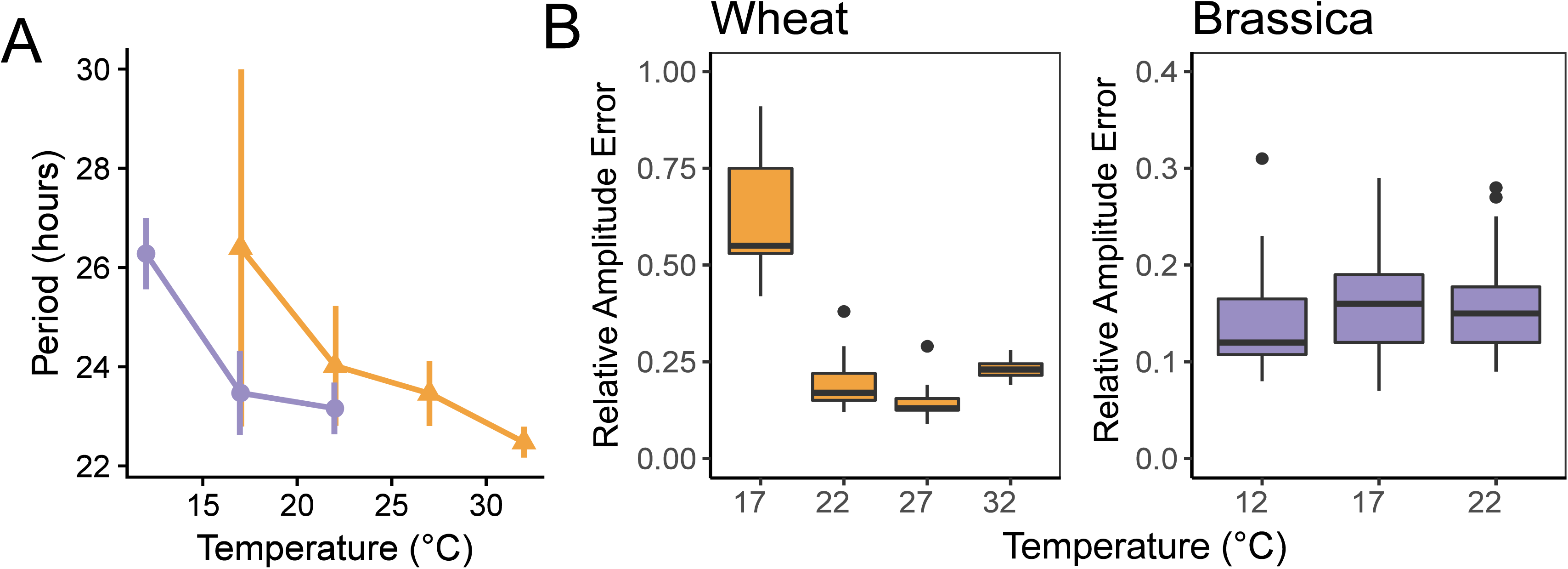
Increasing temperature causes a shortening of period and effects rhythm robustness. Wheat and *Brassica* seedlings were entrained for 4 days in L:D at the temperature being assessed before imaging. Each temperature point represents a separate imaging experiment. Period means decrease with increasing temperatures as is shown in A for *Brassica* (purple circles) or wheat (orange triangles). Error bars represent standard deviation. Box plots show the effect of temperature on RAE for wheat (orange) or *Brassica* (purple) (B). Period and RAE were estimated by FFT NLLS, BAMP dtr, 24-120h cut-off. Wheat: 17°C (N=17/23); 22°C (N=32/32); 27°C (N=11/11); 32°C (N=15/15). *Brassica:* 12°C (N=24/24); 17°C (N=35/35); 22°C (N=30/30). An additional preliminary experiment consistent with these observations can be seen in Supplementary S13. Significance codes: \*\*\**p*<0.001.

Across the temperatures tested *Brassica* rhythm robustness remained consistent; all samples were returned from FFT-NLLS and RAE, period CV and Period error were similar (Figure 5B, Supplementary Materials S6). We recommend 22°C for DF using *Brassica* as it had the lowest period CV of 2.24%.

### An optimized DF method can be used in circadian analysis for crops

To see whether the optimized method could be used to investigate circadian differences between cultivars of the same species, we looked at circadian rhythms from seven *T. aestivum* cultivars and three *B. napus* cultivars. For the *B. napus* lines, seeds were obtained from 3 different harvest years to see whether period was constant between batches. The optimized imaging parameters we used for these elite cultivars is outlined in Table 1.

**Table 1.**
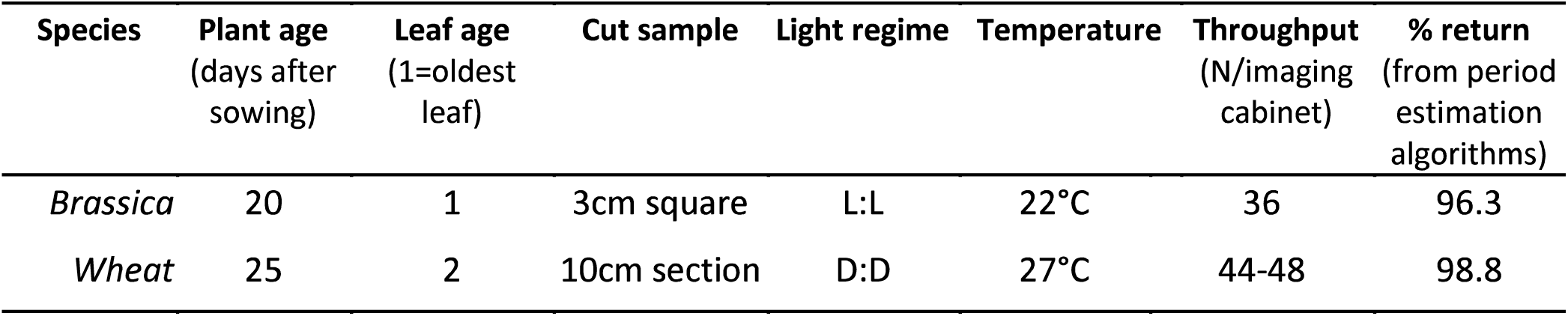
Optimised DF method for circadian phenotyping of *Brassica* and wheat leaves

| Species | Plant age<br>(days after<br>sowing) | Leaf age<br>(1=oldest<br>leaf) | Cut sample | Light regime | Temperature | Throughput<br>(N/imaging<br>cabinet) | % return<br>(from period<br>estimation<br>algorithms) |
| --- | --- | --- | --- | --- | --- | --- | --- |
| *Brassica* | 20 | 1 | 3cm square | L:L | 22°C | 36 | 96.3 |
| *Wheat* | 25 | 2 | 10cm section | D:D | 27°C | 44-48 | 98.8 |

There was significant variation in the periods of the wheat lines tested as shown in Figure 6A (F(6,152)=9.81, p<0.001, one-way ANOVA). A Tukey HSD test showed that Paragon (mean=23.48h, SD=0.54) and Norin 61 (mean=23.50h, SD=1.30) both have longer periods than Chinese Spring (mean=22.70h, SD=0.40), Claire (mean=22.48, SD=0.54) and Robigus (mean=22.32h, SD=0.34) α=0.01).

**Figure 6.**
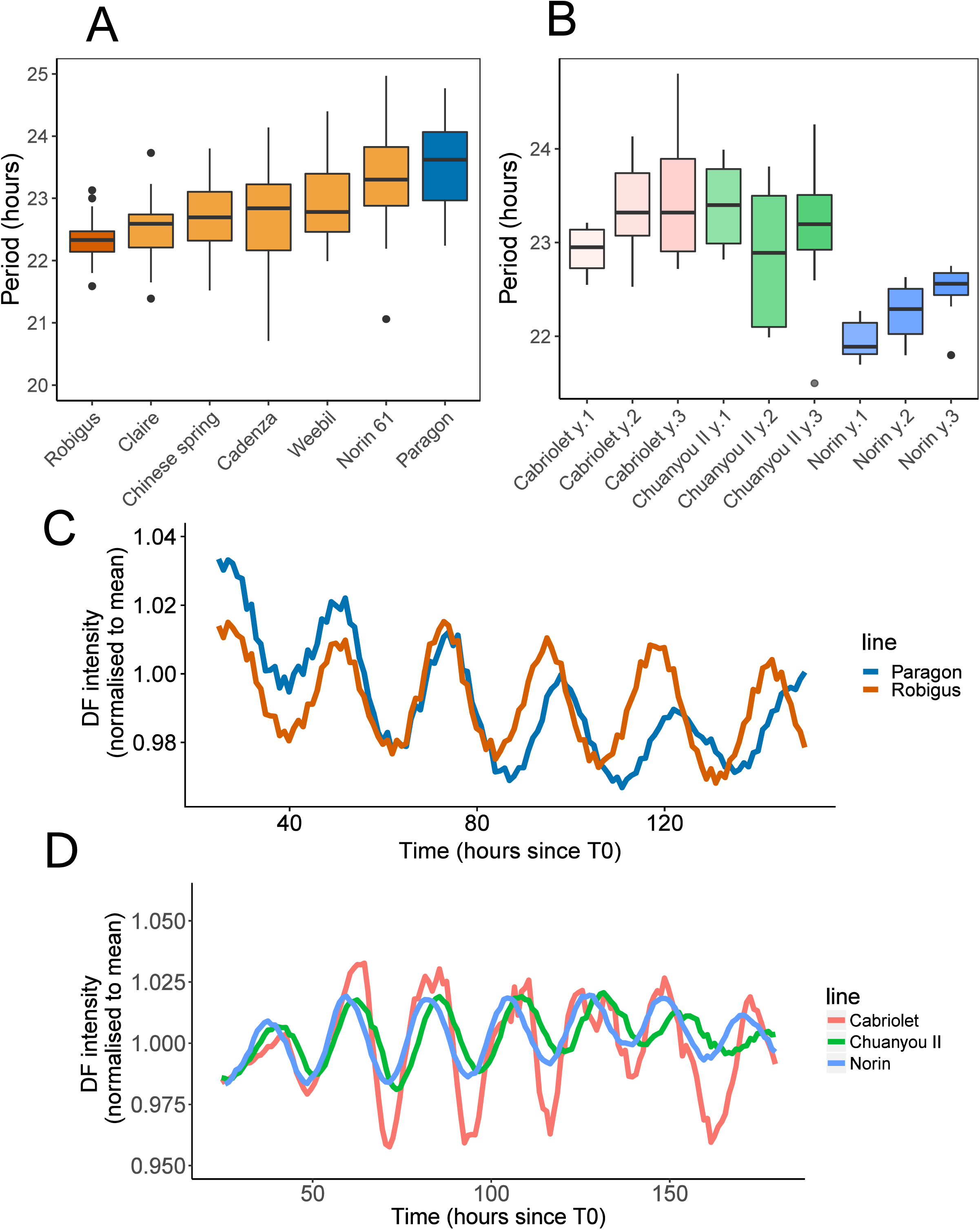
DF can be used to measure period differences between elite cultivars in *Brassica* and wheat. 10cm sections from the second leaf of 25 day old wheat seedlings were imaged under D:D at 27°C. 3cm square sections from the first leaf of 21 day old *Brassica* seedlings were imaged under L:L at 22°C. Period boxplots based on the DF oscillations from different cultivars are shown for wheat (A) and *Brassica* (B). The three replicates in the *Brassica* data represent different seed batches. Period values were estimated using FFT NLLS, BAMP dtr, 24-120h cut-off window. BAMP de-trended DF data was normalised to the mean DF intensity across all cultivars and plotted against time in hours after dawn (C and D). *Brassica* data represents results from two cabinets normalised for between-cabinet effects. Wheat data represents results from two identical experiments with two imaging cabinets run in parallel as technical replicates and normalised for the between-cabinet and experimental-run effects. Wheat: Robigus (N=25/25), Cadenza (N=12/12), Chinese Spring (N=24/25), Claire (N=27/27), Weebil (N=22/24), Paragon (N=24/4), Norin-61 (N=25/25). *Brassica:* for each seed batch of Cabriolet, Chuanyou II, Norin (N=8/8). *Brassica* data is consistent with previous experiments shown in supplementary materials S14. Significance codes: for wheat cultivars (A),\*\*\**p*<0.001, \*\**p*<0.01 relative to either *a*=Norin 61 or *b*=Paragon. For brassica cultivars (B), significantly different periods (*p*<0.05) are labelled relative to *a*=Norin Y1, *b*=Norin Y2 or *c*=Norin Y3.

Figure 6A shows the variation in period across three *Brassica* lines taken from three seed batches. We conducted a two-way analysis of variance to compare cultivar ID and seed batch effects as well as the interaction between the two factors. The cultivar ID was found to have a significant effect on period (F(2,60)=25.47, *p*<0.001) but batch year did not significantly account for any variation in period either as a main effect (F(2,60)=1.73, *p*>0.1) or as an interaction with the cultivar ID (F(4,60)=2.27, *p*=0.72). This suggests that the observed differences in period are due to heritable genetic differences. The *Brassica* cultivar Norin had the shortest overall period of 22.29h (SD 0.34); shorter than either Cabriolet (23.32h, SD=0.57) or Chuanyou II (23.18h, SD=0.72) (*p*<0.001, Tukey HSD).

DF oscillations in both *Brassica* and wheat remained rhythmic throughout the experiment allowing confident period estimation over 4 days (24-120h following T_0_). The average DF oscillations for the two most divergent wheat lines is shown in Figure 6C; the other lines have been omitted for clarity. DF expression from all three years was averaged to make the oscillation plots for the *Brassica* lines as shown in Figure 6D. The percentage of DF rhythms returned from period estimation was high for both *Brassica* (96.3%) and wheat (98.8%) proving that the method is both efficient and reliable.

## Discussion

Manipulating the circadian clock has potential for influencing crop productivity, efficiency and resilience; however research has been hindered by the lack of high-throughput circadian protocols which can be reliably applied to crop plants. Transcriptional assays, luciferase constructs and fluorescent markers have been used to investigate circadian rhythms in tobacco(13), tomato(39), potato(40,41), *Brassica rapa*(42), rice(43-45), barley(46) and wheat(47). However, these approaches are either manually intensive, technologically expensive or require genetic modification to systematically investigate each component and so are low throughput. We have optimized a delayed fluorescence imaging method for reliable circadian phenotyping of either *Brassica* or wheat seedlings. *Brassica napus* (AACC) and *Triticum aestivum* (AABBDD) are both recent polyploids still undergoing genomic rearrangements. The contribution of each genome to clock function remains to be investigated. *B. napus* is a dicot recently diverged from *Arabidopsis*(48) and so is likely to have clock homologs with similar functions. *T. aestivum* is a monocot with an incompletely understood clock mechanism(49). These species therefore provide interesting insights into two genetically diverse families. Several differences between the function of these clocks have been exposed through the factors examined in this paper. The opposing robustness of clocks under D:D or L:L and the fact that DF rhythms peak with distinct phases under each condition is indicative of diverging networks underlying circadian control of each species. Lower temperatures (17°C) also seem to have a detrimental effect on the robustness of the clock in *T. aestivum* but not *B. napus*, suggesting that temperature may be a stronger zeitgeber for wheat than for *Brassica* within this temperature range. The DF rhythms in both *T. aestivum* and *B. napus* have reduced temperature compensation compared to those reported for leaf movement in *Arabidopsis*(36,37). However, it is important to recognize that our rhythms were measured in dissected sections of leaves and may not be truly analogous to rhythms from whole *Arabidopsis* individuals. The difference between intact and excised leaves has been previously reported in Hall et al. 2001(22). Our analysis of the homogeneity of periods across a single leaf also revealed variability of period robustness across the axis of wheat leaves but relatively little variation across *Brassica* leaves.

In this study, we have shown that in both *Brassica* and wheat there is a strong interaction between circadian period and age due to both systemic aging and leaf-specific developmental aging. Previous research in *Arabidopsis* has asked whether the onset of senescence is a result of a faster running clock or vice versa(28,50). Our results suggest that the acceleration of the clock occurs in very young plants before senescence phase, raising the possibility that the clock could be artificially manipulated to moderate senescence and control timing of peak productivity in crops.

Natural variation of circadian phenotypes has been previously demonstrated in wild *Arabidopsis* accessions (35,36,51) revealing a selection pressure for circadian traits specific to different ecological settings. The extent to which circadian fitness has been selected-for in modern crop plants has not yet been investigated. Application of our optimized protocol in this study demonstrates that diverging rhythms are present within elite cultivars of the same species. This variation in circadian period suggests that some level of circadian diversity exists, but the question remains as to whether each cultivar is currently optimized to enhance individual plant fitness. Crop plants with ‘optimized circadian clocks’ may have the capacity to improve yield, efficiency and resilience potentially overlooked by traditional plant breeding methods.

## Conclusions

In this study, we investigated several important factors influencing circadian rhythms in *Brassica napus* and *Triticum aestivum* and reveal intriguing differences between the two crops. We provide an optimized DF methodology which can be reliably used for high-throughput measurement of circadian rhythms. This research highlights the considerable plasticity of the circadian clock under free-running conditions. It is our hope that these results may inform future research by showing the extent to which controllable variables can affect period estimation and how these may differ depending on the model species being studied.

## List of Abbreviations

DF: Delayed Fluorescence
FFT-NLLS: Fast Fourier Transform Non-Linear Least Squares
RAE: Relative Amplitude Error
PSII: Photosystem II
L:L: Constant light
L:D: Light-dark cycles
D:D: Constant Dark
BnDFFS: *Brassica napus* Diversity Fixed Foundation Set
ZT: Zeitgeber time
BAMP: Baseline and amplitude
CV: coefficient of variation

## Method

### Plant material and growth conditions

*Brassica* seedlings used were from the winter varieties Cabriolet and Norin and the semiwinter variety Chuanyou II from the OREGIN *Brassica napus* Diversity Fixed Foundation Set (BnDFFS)(52). Wheat seedlings used were all hexaploid elite cultivars ordered from the Genome Resource Unit (John Innes Centre) (Supplementary Materials S8).

*Brassica* plants were grown in Levington’s F2 mix in FP11 pots, spaced 5 plants to a pot. They were grown in controlled greenhouse conditions, (16:8h L:D at 22:20°C). After 17 days, plants were transferred to a plant growth chamber set at 12:12 L:D cycle at 22°C under approximately 200μmol m^−2^ s^−1^ white light for 4 days entrainment (light spectra can be seen in Supplementary Materials S9).

Wheat plants were imbibed at 4°C for 6 days before being planted in Petersfield cereal mix in FP9 pots, spaced two to a pot. They were then grown in controlled greenhouse conditions (16:8h L:D 17:12°C). After 21 days plants were transferred to a plant growth chamber set at the cabinet conditions above. For temperature experiments, plants were entrained at the temperatures in which they would be imaged.

### Image acquisition-standard conditions

Leaves were removed just after entrainment dawn and placed face up onto 24cm square petri dishes (Stratlab LTD, cat no. 163-PB-007) containing 0.5% water agar (Sigma-Aldrich, SKU A1296). Unless otherwise stated, 3cm squares were cut from the second true leaf of 21 day old *B. napus* seedlings. A segment of 10cm was taken from the second leaf of the main tiller of 25 day old *T. aestivum* seedlings, beginning 5cm down from the tip. Plates were secured with masking tape around the periphery.

The imaging set-up is adapted from that described by Southern et al(53). A set-up schematic can be seen in Supplementary Materials S10. We use Lumo Reteiga CCD cameras (Qlmaging, Canada), which we have found to have comparable image quality to the Orea II (Hamamatsu Photonics, Japan) without the need to run a water-cooling pump. Cameras were fitted with a Xenon 0.95/25mm lens (Schneider-Kreuznach, Germany).

A custom built 25×25 red/blue LED rig (approx. 60μmol m^−2^ s^−1^) was controlled by μManager software (v1.4.19, Open Imaging) through an Arduino Uno microcontroller board(54). LED spectra for cabinets can be viewed in Supplementary Materials S9. μManager was used to configure both the supplied camera driver software (PVCam v3.7.1.0) and program the Arduino after installing the firmware source code available online (55)).

Both camera and LEDs were housed in a temperature controlled growth cabinet (Sanyo MIR-553) in a dark room. The temperature was set to 22°C unless otherwise specified (changed for the temperature experiments (Figure 5) and for the wheat cultivar experiment (Figure 6)). Camera properties were kept the same in each experiment (Binning=4, Gain=1, Readout-Rate=0.650195MHz 16 bit) and camera exposure was initiated 500ms after the lights were turned off. A ‘L:L’ script refers to a regime of 59min of light followed by a 1 minute exposure in the dark. A ‘D:D’ script refers to 54min of darkness followed by 5min light and then the 1min exposure. BeanShell scripts run by μManager have been adapted from scripts used previously(56) and are available to view as Additional files 15 and 16. Wheat imaging used the D:D script and *Brassica* imaging used the L:L script with the exception of experiments in Figure 4.

### Processing in FIJI and BioDare2 parameters

Image stacks were imported into FIJI(57) and regions of interest were selected. Measurements for integrated density were taken for these regions across the stack using the Multi-measure plugin. Each region was then labeled in Excel and an offset time series added. The ‘offset time’ is the difference between the time of the first image (T1) and entrainment dawn (ZT) in decimal hours. Data can then be uploaded to BioDare2 as described online(58,59). BioDare2 is an open-access web tool for analyzing timeseries data and predicting circadian parameters. For our data we found that Baseline and amplitude (BAMP) de-trending was most appropriate but recommend visual inspection of the detrending methods available to find the least intrusive method which removes any baseline trends. Period estimation was done using the Fast Fourier Transform Non-Linear Least Squares (FFT-NLLS) algorithm(60) on a data window of 24-120h with expected periods set to 18-34h. Manual inspection of resulting periods ensured that all arrhythmic traces were excluded from further analysis.

### Rhythm Robustness analysis

We summarized rhythm robustness metrics based on several BioDare2 outputs.’% returned’ is the number of samples for which periods could be estimated out of the number of samples originally imaged. The RAE (relative amplitude error) is the ratio of amplitude error to amplitude and represents amplitude robustness. A RAE of 1 indicates the most irregular waveform which can still be classified as rythmic whereas a RAE of 0 indicates a perfect sine wave with no amplitude error. The period coefficient of variation (CV) is the standard deviation of period estimates adjusted for the mean period and represents between sample variation(58,61). Period error is the extent to which the period estimate could vary and still give a good fit to the model. Error scores close to 0 indicate a tight fit of the model to the observed data and a high within sample period robustness. See Supplementary data Sl-6 for statistic tables and further descriptions.

### Normalization for experimental effects

After circadian parameters were estimated in BioDare2, data was normalized to account for the following random experimental effects. For the wheat plant age and leaf age experiments (Figure 1), *Brassica* plant-leaf age experiments (Figure 2) and the *Brassica* cultivar experiments (Figure 6B) samples were split between two imaging cabinets run in parallel in a single experiment. The predicted parameters (e.g. period) for each sample from the two cabinets were adjusted so that the cabinet means were then equivalent. This was achieved by dividing the cabinet means by the overall mean to get an adjustment factor for each cabinet and then dividing each individual value by that factor to get a cabinet-normalized value. For the cutting data (Figure 3), the experiments were replicated in two separate imaging weeks and then adjusted for the between-experiment effects. For the wheat cultivar experiments (Figure 6A) data was obtained from two cabinets over two separate experiments and was normalized for both effects in a similar way. The light regime (Figure 4) and the temperature experiments (Figure 5) measured each variable in one cabinet at a time and therefore did not require any normalization. The conclusions from these experiments is consistent with preliminary experiments presented in Supplementary Materials S11-14.

Statistical analysis was carried out in RStudio v1.1.423 using aov and t.test functions fit with an appropriate linear model in the format specified in the Results.

### Declarations

#### Ethics approval and consent to participate

Not applicable

#### Consent for publication

Not applicable

#### Availability of data and materials

The datasets generated during the current study are available as additional files and supplementary materials in the online version of this article. Raw image files are available from the corresponding author on reasonable request.

### Authors’ contributions

This project was conceptualized by HR and AH. HR designed and conducted experiments, carried out data processing and analysis and wrote the manuscript with contributions from AH. All authors read and approved the final manuscript.

### Abbreviations

Hannah Rees (HR), Susan Duncan (SD), Peter Gould (PG), Rachel Wells (RW), Mark Greenwood (MG), Thomas Brabbs (TB), Anthony Hall (AH)

