## Supplementary for "A high-throughput delayed fluorescence method reveals underlying differences in the control of circadian rhythms in *Triticum aestivum* and *Brassica napus*"

| **Explanation of statistics used to assess robustness**  The tested variable with the most robust statistic is shown in bold in a grey highlighted box | |
| --- | --- |
| % returned from FFTNLLS | Percentage of samples for which the algorithm could calculate a period estimate and define as ‘rhythmic’ |
| RAE ratio average | Average ratio of amplitude error to amplitude. Represents amplitude robustness. |
| Period CV(%) | The coefficient of variation for all period estimates. Represents between sample period robustness. |
| Average of Period Err threshold | Average Period error-threshold for which the period estimate could vary and still give a good fit to the model (FFT-NLLS). Represents within sample period robustness. |

Rhythm robustness testing

**S1. Rhythm robustness table (wheat age Exp.1)**

|  | % returned from FFTNLLS | RAE ratio average | Period CV(%) | Average of Period Err threshold |
| --- | --- | --- | --- | --- |
| 18 days after sowing | **100** | 0.17 | 2.10 | 0.58 |
| 25 days after sowing | **100** | 0.16 | **1.52** | 0.51 |
| 32 days after sowing | 96.15 | **0.15** | 1.98 | **0.43** |
| 39 days after sowing | 82.61 | 0.22 | 3.23 | 0.67 |

|  | % returned from FFTNLLS | RAE ratio average | Period CV(%) | Average of Period Err threshold |
| --- | --- | --- | --- | --- |
| Leaf 1 | **100** | 0.22 | 3.16 | 0.65 |
| Leaf 2 | **100** | **0.18** | 5.77 | **0.5** |
| Leaf 3 | **100** | 0.23 | 5.25 | 0.66 |
| Leaf 4 | 95.45 | 0.22 | **1.81** | 0.68 |

**S2. Rhythm robustness table (wheat leaf Exp.1)**

**S3. Rhythm robustness table (*Brassica* leaf variation by plant age Exp.2)**

|  | % returned from FFTNLLS | RAE ratio average | Period CV(%) | Average of Period Err threshold |
| --- | --- | --- | --- | --- |
| Age_20_Lea f_1 | **100.00** | **0.15** | 1.89 | **0.47** |
| Age_20_Lea f_3 | **100.00** | 0.19 | **1.68** | 0.72 |
| Age_20_Lea f_5 | **100.00** | 0.16 | 4.18 | 0.74 |
| Age_25_Lea f_1 | 66.67 | 0.38 | 13.44 | 1.25 |
| Age_25_Lea f_3 | **100.00** | 0.28 | 3.06 | 1.05 |
| Age_25_Lea f_5 | **100.00** | 0.17 | 2.67 | 0.57 |
| Age_30_Lea f_1 | 33.33 | 0.37 | 2.01 | 1.34 |
| Age_30_Lea f_3 | 83.33 | 0.26 | 5.70 | 0.73 |
| Age_30_Lea f_5 | **100.00** | 0.25 | 2.41 | 0.72 |
| Age_35_Lea f_1 | **100.00** | 0.27 | 5.04 | 1.05 |
| Age_35_Lea f_3 | **100.00** | 0.37 | 4.49 | 1.10 |
| Age_35_Lea f_5 | **100.00** | 0.27 | 2.86 | 0.69 |

**S4. Rhythm robustness table (wheat and *Brassica* cutting samples Exp.3)**

|  | % returned from FFTNLLS | RAE ratio average | Period CV(%) | Average of Period Err threshold |
| --- | --- | --- | --- | --- |
| **WHEAT** | | | | |
| Whole | **100** | 0.16 | 1.99 | 0.47 |
| 10cm | **100** | **0.15** | **1.89** | **0.41** |
| 4cm top | 95.24 | 0.17 | 2.12 | 0.45 |
| 4cm bottom | **100** | 0.16 | 2.72 | 0.45 |
| **BRASSICA** | | | | |
| Whole | **100** | **0.16** | **1.89** | **0.54** |
| Square | **100** | **0.16** | 2.10 | **0.54** |
| Quarter | 98.68 | 0.17 | 2.60 | **0.54** |

**S5. Rhythm robustness table (D:D V L:L Exp.4)**

|  | % returned | RAE average | Period CV | Average of Period Err |
| --- | --- | --- | --- | --- |
| *Brassica* DD | 66.67 | 0.53 | 11.80 | 1.62 |
| *Brassica* LL | **100.00** | **0.13** | **1.34** | **0.41** |
| Wheat DD | **100.00** | **0.20** | **2.25** | **0.55** |
| Wheat LL | 95.83 | 0.38 | 13.53 | 1.17 |

**S6. Rhythm robustness table (Temperature Exp.5)**

|  | % returned from FFTNLLS | RAE ratio average | Period CV(%) | Average of Period Err threshold |
| --- | --- | --- | --- | --- |
| **BRASSICA** | | | | |
| 12 | **100.00** | **0.14** | 2.72 | 0.55 |
| 17 | **100.00** | 0.16 | 3.61 | **0.50** |
| 22 | **100.00** | 0.16 | **2.24** | 0.52 |
| **WHEAT** | | | | |
| 17 | 73.91 | 0.64 | 13.63 | 2.69 |
| 22 | **100.00** | 0.19 | 5.01 | 0.57 |
| 27 | **100.00** | **0.15** | 2.78 | **0.48** |
| 32 | **100.00** | 0.23 | **1.38** | 0.74 |

**S7. Calculation of Q10 temperature coefficient values**

$Q_{10}=\left( \frac{R2}{R1} \right)^{10/(T2-T1)}$

Where:

R=Period as a rate: 1 oscillation/time (hours). E.g: 1/24=0.042 oscillations/hour

T=Temperature °C

R1 is the rate at T1. R2 is the rate at T2.

**S8. Wheat cultivar Genome Resource Unit information:**

| **Line** | **Collection** | **Origin Country** | **Sowing Season** | **Type** |
| --- | --- | --- | --- | --- |
| Norin-61 | BBSRC_Wheat | Japan | Winter | Breeders line |
| Robigus | BBSRC_Wheat | UK | Winter | Advanced/improved cultivar |
| Claire | BBSRC_Wheat | UK | Winter | Advanced/improved cultivar |
| Paragon | BBSRC_Wheat | UK | Spring | Advanced/improved cultivar |
| Cadenza | BBSRC_Wheat | UK | Spring | Founder stock/base population |
| Weebil | WBCDB | Mexico | Spring | Founder stock/base population |
| Chinese Spring L42 | WBCDB | China | Spring | Founder stock/base population |

**S9. Spectral readings of light inside entrainment and imaging cabinets:**

LED spectra inside imaging cabinets (Lumo 1 and Lumo 2)

% saturation

| Wavelength (nm) |  |  |
| --- | --- | --- |
| 380 | 0.069908 | 0.074552 |
| 381 | 0.069191 | 0.074106 |
| 382 | 0.068523 | 0.07356 |
| 383 | 0.067941 | 0.072836 |
| 384 | 0.066841 | 0.071662 |
| 385 | 0.0652 | 0.070019 |
| 386 | 0.063267 | 0.067996 |
| 387 | 0.061151 | 0.065737 |
| 388 | 0.059199 | 0.063738 |
| 389 | 0.057303 | 0.061829 |
| 390 | 0.055928 | 0.060369 |
| 391 | 0.05463 | 0.058975 |
| 392 | 0.05382 | 0.05807 |
| 393 | 0.053068 | 0.057185 |
| 394 | 0.052693 | 0.05644 |
| 395 | 0.052393 | 0.055847 |
| 396 | 0.052301 | 0.055685 |
| 397 | 0.052139 | 0.055423 |
| 398 | 0.051869 | 0.055002 |
| 399 | 0.051679 | 0.054969 |
| 400 | 0.051563 | 0.055292 |
| 401 | 0.051525 | 0.055394 |
| 402 | 0.051528 | 0.055377 |
| 403 | 0.051771 | 0.055277 |
| 404 | 0.05208 | 0.055154 |
| 405 | 0.051356 | 0.05481 |
| 406 | 0.050588 | 0.054456 |
| 407 | 0.050954 | 0.054318 |
| 408 | 0.05109 | 0.054105 |
| 409 | 0.050163 | 0.053549 |
| 410 | 0.049415 | 0.053015 |
| 411 | 0.049061 | 0.052532 |
| 412 | 0.048651 | 0.051966 |
| 413 | 0.048171 | 0.051298 |
| 414 | 0.047716 | 0.050747 |
| 415 | 0.047279 | 0.050282 |
| 416 | 0.046903 | 0.049947 |
| 417 | 0.046553 | 0.049664 |
| 418 | 0.046885 | 0.049955 |
| 419 | 0.047338 | 0.050347 |
| 420 | 0.048173 | 0.051321 |
| 421 | 0.049111 | 0.052357 |
| 422 | 0.050738 | 0.053776 |
| 423 | 0.052425 | 0.055291 |
| 424 | 0.054281 | 0.057078 |
| 425 | 0.056483 | 0.059391 |
| 426 | 0.059206 | 0.062499 |
| 427 | 0.062764 | 0.065947 |
| 428 | 0.067041 | 0.069689 |
| 429 | 0.071882 | 0.074407 |
| 430 | 0.076992 | 0.079592 |
| 431 | 0.08384 | 0.086214 |
| 432 | 0.091077 | 0.093158 |
| 433 | 0.100397 | 0.102343 |
| 434 | 0.110062 | 0.111845 |
| 435 | 0.122008 | 0.123322 |
| 436 | 0.134875 | 0.13559 |
| 437 | 0.150621 | 0.150331 |
| 438 | 0.167922 | 0.166817 |
| 439 | 0.187701 | 0.186083 |
| 440 | 0.210719 | 0.208002 |
| 441 | 0.236611 | 0.232276 |
| 442 | 0.266593 | 0.260723 |
| 443 | 0.298544 | 0.291179 |
| 444 | 0.337253 | 0.327854 |
| 445 | 0.377433 | 0.365884 |
| 446 | 0.42469 | 0.411217 |
| 447 | 0.472997 | 0.457668 |
| 448 | 0.527531 | 0.5108 |
| 449 | 0.582846 | 0.564831 |
| 450 | 0.64037 | 0.621412 |
| 451 | 0.697415 | 0.677864 |
| 452 | 0.753759 | 0.734127 |
| 453 | 0.806697 | 0.787595 |
| 454 | 0.856863 | 0.838788 |
| 455 | 0.899027 | 0.882903 |
| 456 | 0.937732 | 0.923959 |
| 457 | 0.96403 | 0.953439 |
| 458 | 0.98807 | 0.98081 |
| 459 | 0.995075 | 0.991469 |
| 460 | 1 | 1 |
| 461 | 0.993142 | 0.996455 |
| 462 | 0.98168 | 0.988062 |
| 463 | 0.959229 | 0.968097 |
| 464 | 0.931934 | 0.942542 |
| 465 | 0.898649 | 0.910074 |
| 466 | 0.861708 | 0.8738 |
| 467 | 0.82233 | 0.834989 |
| 468 | 0.782161 | 0.794159 |
| 469 | 0.741733 | 0.75267 |
| 470 | 0.700929 | 0.710878 |
| 471 | 0.660197 | 0.669161 |
| 472 | 0.621057 | 0.628945 |
| 473 | 0.582725 | 0.5896 |
| 474 | 0.547215 | 0.553294 |
| 475 | 0.512505 | 0.51779 |
| 476 | 0.479094 | 0.483587 |
| 477 | 0.447595 | 0.451268 |
| 478 | 0.417718 | 0.420549 |
| 479 | 0.389528 | 0.391882 |
| 480 | 0.362055 | 0.364088 |
| 481 | 0.336681 | 0.338525 |
| 482 | 0.311722 | 0.313407 |
| 483 | 0.289822 | 0.291639 |
| 484 | 0.268506 | 0.270387 |
| 485 | 0.249659 | 0.251316 |
| 486 | 0.231615 | 0.233035 |
| 487 | 0.215019 | 0.216178 |
| 488 | 0.199669 | 0.200696 |
| 489 | 0.185451 | 0.186463 |
| 490 | 0.172936 | 0.173879 |
| 491 | 0.161178 | 0.162026 |
| 492 | 0.151022 | 0.151917 |
| 493 | 0.141168 | 0.142137 |
| 494 | 0.132691 | 0.133841 |
| 495 | 0.124513 | 0.125782 |
| 496 | 0.117498 | 0.118653 |
| 497 | 0.110768 | 0.111911 |
| 498 | 0.104507 | 0.105809 |
| 499 | 0.098803 | 0.100223 |
| 500 | 0.093551 | 0.095055 |
| 501 | 0.088511 | 0.090184 |
| 502 | 0.08355 | 0.085425 |
| 503 | 0.079403 | 0.081265 |
| 504 | 0.075377 | 0.077188 |
| 505 | 0.071754 | 0.073351 |
| 506 | 0.068223 | 0.069755 |
| 507 | 0.064953 | 0.066835 |
| 508 | 0.062076 | 0.064051 |
| 509 | 0.059688 | 0.061436 |
| 510 | 0.057546 | 0.059437 |
| 511 | 0.055549 | 0.057802 |
| 512 | 0.053715 | 0.05589 |
| 513 | 0.051919 | 0.053915 |
| 514 | 0.050555 | 0.052531 |
| 515 | 0.049154 | 0.051109 |
| 516 | 0.047585 | 0.049521 |
| 517 | 0.046143 | 0.048105 |
| 518 | 0.044912 | 0.046977 |
| 519 | 0.043689 | 0.045755 |
| 520 | 0.04247 | 0.04446 |
| 521 | 0.0417 | 0.043702 |
| 522 | 0.041074 | 0.043119 |
| 523 | 0.040484 | 0.042509 |
| 524 | 0.039889 | 0.041942 |
| 525 | 0.039245 | 0.04166 |
| 526 | 0.038632 | 0.041186 |
| 527 | 0.038089 | 0.040286 |
| 528 | 0.037641 | 0.039711 |
| 529 | 0.03729 | 0.039472 |
| 530 | 0.036718 | 0.03906 |
| 531 | 0.03604 | 0.038565 |
| 532 | 0.035893 | 0.038199 |
| 533 | 0.035815 | 0.037855 |
| 534 | 0.035384 | 0.037544 |
| 535 | 0.035122 | 0.037327 |
| 536 | 0.035412 | 0.037421 |
| 537 | 0.03541 | 0.037426 |
| 538 | 0.035011 | 0.03731 |
| 539 | 0.034836 | 0.037235 |
| 540 | 0.034802 | 0.037185 |
| 541 | 0.034699 | 0.037019 |
| 542 | 0.03458 | 0.036825 |
| 543 | 0.034558 | 0.036692 |
| 544 | 0.034504 | 0.036584 |
| 545 | 0.034317 | 0.036577 |
| 546 | 0.034299 | 0.036651 |
| 547 | 0.034539 | 0.036852 |
| 548 | 0.034642 | 0.037022 |
| 549 | 0.034652 | 0.037172 |
| 550 | 0.034699 | 0.037162 |
| 551 | 0.034755 | 0.037111 |
| 552 | 0.034688 | 0.037095 |
| 553 | 0.034638 | 0.037073 |
| 554 | 0.034656 | 0.037022 |
| 555 | 0.034543 | 0.036911 |
| 556 | 0.034236 | 0.036711 |
| 557 | 0.034035 | 0.036562 |
| 558 | 0.033899 | 0.036446 |
| 559 | 0.033977 | 0.036431 |
| 560 | 0.034106 | 0.036444 |
| 561 | 0.03433 | 0.03667 |
| 562 | 0.034457 | 0.036793 |
| 563 | 0.034275 | 0.0366 |
| 564 | 0.034187 | 0.036467 |
| 565 | 0.034208 | 0.036406 |
| 566 | 0.034143 | 0.036432 |
| 567 | 0.034034 | 0.0365 |
| 568 | 0.034137 | 0.036617 |
| 569 | 0.034233 | 0.036718 |
| 570 | 0.034052 | 0.036626 |
| 571 | 0.033978 | 0.036553 |
| 572 | 0.034121 | 0.03652 |
| 573 | 0.034217 | 0.036651 |
| 574 | 0.034276 | 0.036915 |
| 575 | 0.034453 | 0.037026 |
| 576 | 0.034664 | 0.037093 |
| 577 | 0.034679 | 0.036979 |
| 578 | 0.034729 | 0.036999 |
| 579 | 0.034894 | 0.037467 |

Fluorescent tube spectra inside entrainmnet cabinets (5 bar)

% saturation

| Wavelength (nm) |  |  |
| --- | --- | --- |
| 380 | 0.099274 | 0.084456 |
| 381 | 0.097886 | 0.08323 |
| 382 | 0.096663 | 0.081979 |
| 383 | 0.095733 | 0.080682 |
| 384 | 0.094065 | 0.079059 |
| 385 | 0.091628 | 0.077096 |
| 386 | 0.089129 | 0.075198 |
| 387 | 0.086592 | 0.07334 |
| 388 | 0.084323 | 0.071577 |
| 389 | 0.082146 | 0.069845 |
| 390 | 0.080189 | 0.068184 |
| 391 | 0.078267 | 0.066535 |
| 392 | 0.077237 | 0.06567 |
| 393 | 0.076236 | 0.064789 |
| 394 | 0.075426 | 0.063807 |
| 395 | 0.075264 | 0.063675 |
| 396 | 0.076944 | 0.065959 |
| 397 | 0.083351 | 0.073005 |
| 398 | 0.097147 | 0.087492 |
| 399 | 0.106372 | 0.096974 |
| 400 | 0.111385 | 0.101844 |
| 401 | 0.114699 | 0.105283 |
| 402 | 0.117103 | 0.107954 |
| 403 | 0.118524 | 0.109732 |
| 404 | 0.119672 | 0.111261 |
| 405 | 0.120195 | 0.111788 |
| 406 | 0.120567 | 0.112122 |
| 407 | 0.119016 | 0.110421 |
| 408 | 0.115223 | 0.106451 |
| 409 | 0.101103 | 0.092025 |
| 410 | 0.09111 | 0.081908 |
| 411 | 0.090242 | 0.081319 |
| 412 | 0.090075 | 0.081393 |
| 413 | 0.090783 | 0.082294 |
| 414 | 0.092561 | 0.084345 |
| 415 | 0.095117 | 0.087232 |
| 416 | 0.097489 | 0.090177 |
| 417 | 0.099787 | 0.093144 |
| 418 | 0.102653 | 0.096766 |
| 419 | 0.105619 | 0.100504 |
| 420 | 0.109396 | 0.104581 |
| 421 | 0.113253 | 0.108768 |
| 422 | 0.117603 | 0.113716 |
| 423 | 0.121956 | 0.118653 |
| 424 | 0.126317 | 0.123561 |
| 425 | 0.13177 | 0.129583 |
| 426 | 0.138875 | 0.13729 |
| 427 | 0.163188 | 0.163819 |
| 428 | 0.202353 | 0.206594 |
| 429 | 0.252175 | 0.26093 |
| 430 | 0.307098 | 0.320798 |
| 431 | 0.327575 | 0.342687 |
| 432 | 0.340313 | 0.356044 |
| 433 | 0.344532 | 0.360738 |
| 434 | 0.348271 | 0.364904 |
| 435 | 0.351077 | 0.367766 |
| 436 | 0.353195 | 0.370067 |
| 437 | 0.353163 | 0.370607 |
| 438 | 0.341127 | 0.35775 |
| 439 | 0.309969 | 0.323549 |
| 440 | 0.268848 | 0.279005 |
| 441 | 0.21889 | 0.225288 |
| 442 | 0.20146 | 0.206558 |
| 443 | 0.199683 | 0.204664 |
| 444 | 0.199792 | 0.20486 |
| 445 | 0.200311 | 0.205512 |
| 446 | 0.200449 | 0.205878 |
| 447 | 0.200619 | 0.206242 |
| 448 | 0.201134 | 0.206663 |
| 449 | 0.201374 | 0.206893 |
| 450 | 0.200831 | 0.206583 |
| 451 | 0.200263 | 0.205981 |
| 452 | 0.199656 | 0.20495 |
| 453 | 0.19867 | 0.203832 |
| 454 | 0.197373 | 0.202645 |
| 455 | 0.196092 | 0.200994 |
| 456 | 0.194817 | 0.199143 |
| 457 | 0.192343 | 0.196776 |
| 458 | 0.18965 | 0.194314 |
| 459 | 0.187711 | 0.191739 |
| 460 | 0.185779 | 0.18921 |
| 461 | 0.183862 | 0.186957 |
| 462 | 0.181726 | 0.184522 |
| 463 | 0.179066 | 0.181653 |
| 464 | 0.176393 | 0.178795 |
| 465 | 0.173705 | 0.175952 |
| 466 | 0.171282 | 0.17306 |
| 467 | 0.169036 | 0.170134 |
| 468 | 0.166261 | 0.167157 |
| 469 | 0.163313 | 0.164163 |
| 470 | 0.160622 | 0.161094 |
| 471 | 0.157962 | 0.158038 |
| 472 | 0.155367 | 0.155279 |
| 473 | 0.153412 | 0.153042 |
| 474 | 0.153685 | 0.152631 |
| 475 | 0.155619 | 0.154285 |
| 476 | 0.160245 | 0.159293 |
| 477 | 0.169543 | 0.168669 |
| 478 | 0.182807 | 0.181753 |
| 479 | 0.200118 | 0.199263 |
| 480 | 0.21915 | 0.218653 |
| 481 | 0.239164 | 0.238307 |
| 482 | 0.259252 | 0.257938 |
| 483 | 0.276467 | 0.275301 |
| 484 | 0.292642 | 0.29165 |
| 485 | 0.304427 | 0.303723 |
| 486 | 0.313559 | 0.313009 |
| 487 | 0.317918 | 0.317277 |
| 488 | 0.317032 | 0.316229 |
| 489 | 0.311382 | 0.310355 |
| 490 | 0.30013 | 0.298711 |
| 491 | 0.286387 | 0.284502 |
| 492 | 0.269311 | 0.266977 |
| 493 | 0.251657 | 0.248882 |
| 494 | 0.232736 | 0.229667 |
| 495 | 0.21404 | 0.210674 |
| 496 | 0.196221 | 0.192541 |
| 497 | 0.179098 | 0.17517 |
| 498 | 0.163122 | 0.159056 |
| 499 | 0.148739 | 0.144381 |
| 500 | 0.135649 | 0.130873 |
| 501 | 0.123524 | 0.11837 |
| 502 | 0.11176 | 0.106243 |
| 503 | 0.103475 | 0.097842 |
| 504 | 0.095818 | 0.090093 |
| 505 | 0.091054 | 0.085202 |
| 506 | 0.086776 | 0.080783 |
| 507 | 0.083848 | 0.077676 |
| 508 | 0.081445 | 0.074998 |
| 509 | 0.079696 | 0.072852 |
| 510 | 0.078065 | 0.071152 |
| 511 | 0.076506 | 0.069714 |
| 512 | 0.075102 | 0.068439 |
| 513 | 0.073735 | 0.067202 |
| 514 | 0.072668 | 0.065976 |
| 515 | 0.071566 | 0.06476 |
| 516 | 0.070308 | 0.063586 |
| 517 | 0.069256 | 0.062499 |
| 518 | 0.068547 | 0.06156 |
| 519 | 0.067734 | 0.06059 |
| 520 | 0.066841 | 0.059596 |
| 521 | 0.066334 | 0.059244 |
| 522 | 0.065951 | 0.0591 |
| 523 | 0.066081 | 0.059314 |
| 524 | 0.066448 | 0.059752 |
| 525 | 0.068191 | 0.061544 |
| 526 | 0.070567 | 0.063945 |
| 527 | 0.074341 | 0.067692 |
| 528 | 0.080146 | 0.073706 |
| 529 | 0.088037 | 0.082053 |
| 530 | 0.10259 | 0.096723 |
| 531 | 0.120337 | 0.114423 |
| 532 | 0.154519 | 0.149059 |
| 533 | 0.193057 | 0.188154 |
| 534 | 0.266253 | 0.262351 |
| 535 | 0.347126 | 0.344364 |
| 536 | 0.453109 | 0.45194 |
| 537 | 0.562693 | 0.562518 |
| 538 | 0.677183 | 0.677185 |
| 539 | 0.776199 | 0.77665 |
| 540 | 0.865489 | 0.866558 |
| 541 | 0.925218 | 0.926377 |
| 542 | 0.977839 | 0.978961 |
| 543 | 0.993338 | 0.993832 |
| 544 | 1 | 1 |
| 545 | 0.97038 | 0.970435 |
| 546 | 0.918932 | 0.9184 |
| 547 | 0.833787 | 0.831678 |
| 548 | 0.732041 | 0.728637 |
| 549 | 0.618848 | 0.614342 |
| 550 | 0.511824 | 0.507142 |
| 551 | 0.406417 | 0.401799 |
| 552 | 0.33786 | 0.333241 |
| 553 | 0.274343 | 0.269684 |
| 554 | 0.230384 | 0.225526 |
| 555 | 0.191299 | 0.186082 |
| 556 | 0.159413 | 0.153598 |
| 557 | 0.133522 | 0.127267 |
| 558 | 0.11145 | 0.104854 |
| 559 | 0.097552 | 0.090717 |
| 560 | 0.085642 | 0.078596 |
| 561 | 0.07979 | 0.072714 |
| 562 | 0.074558 | 0.067398 |
| 563 | 0.071267 | 0.063844 |
| 564 | 0.069049 | 0.061547 |
| 565 | 0.06811 | 0.060747 |
| 566 | 0.068468 | 0.061067 |
| 567 | 0.06946 | 0.061933 |
| 568 | 0.073838 | 0.066127 |
| 569 | 0.079067 | 0.071154 |
| 570 | 0.087826 | 0.079616 |
| 571 | 0.098191 | 0.089793 |
| 572 | 0.111825 | 0.10346 |
| 573 | 0.127524 | 0.119014 |
| 574 | 0.144899 | 0.1361 |
| 575 | 0.164938 | 0.156399 |
| 576 | 0.18575 | 0.177628 |
| 577 | 0.208709 | 0.200849 |
| 578 | 0.23168 | 0.224128 |
| 579 | 0.254696 | 0.247599 |

**S10. Set up of equipment for delayed fluorescence:**

An Arduino USB 2.0 cable connects the computer (1) to an Arduino board (2) controlled by a uManager BSH script. The Arduino sends a signal to a mains relay through two jumper wires (3) which converts power to 12V to turn the LED rig on and off.

The Lumo Reteiga CCD camera is mounted within a temperature regulated Sanyo MIR-553 growth cabinet. A Xenon 0.95/25mm lens is attached to the camera which is focused on a stage carrying the plate to be imaged. The lens fits through a customized hole in the LED rig.

The cabinet and camera are contained within a dark room sprayed matt black and the LEDs used have no afterglow so that total darkness can be created (grey box). The computer and controllers are positioned in the lab for ease of access (yellow box). Data from images taken by the camera are displayed post acquisition and are also automatically saved in a drive specified by the BSH script.

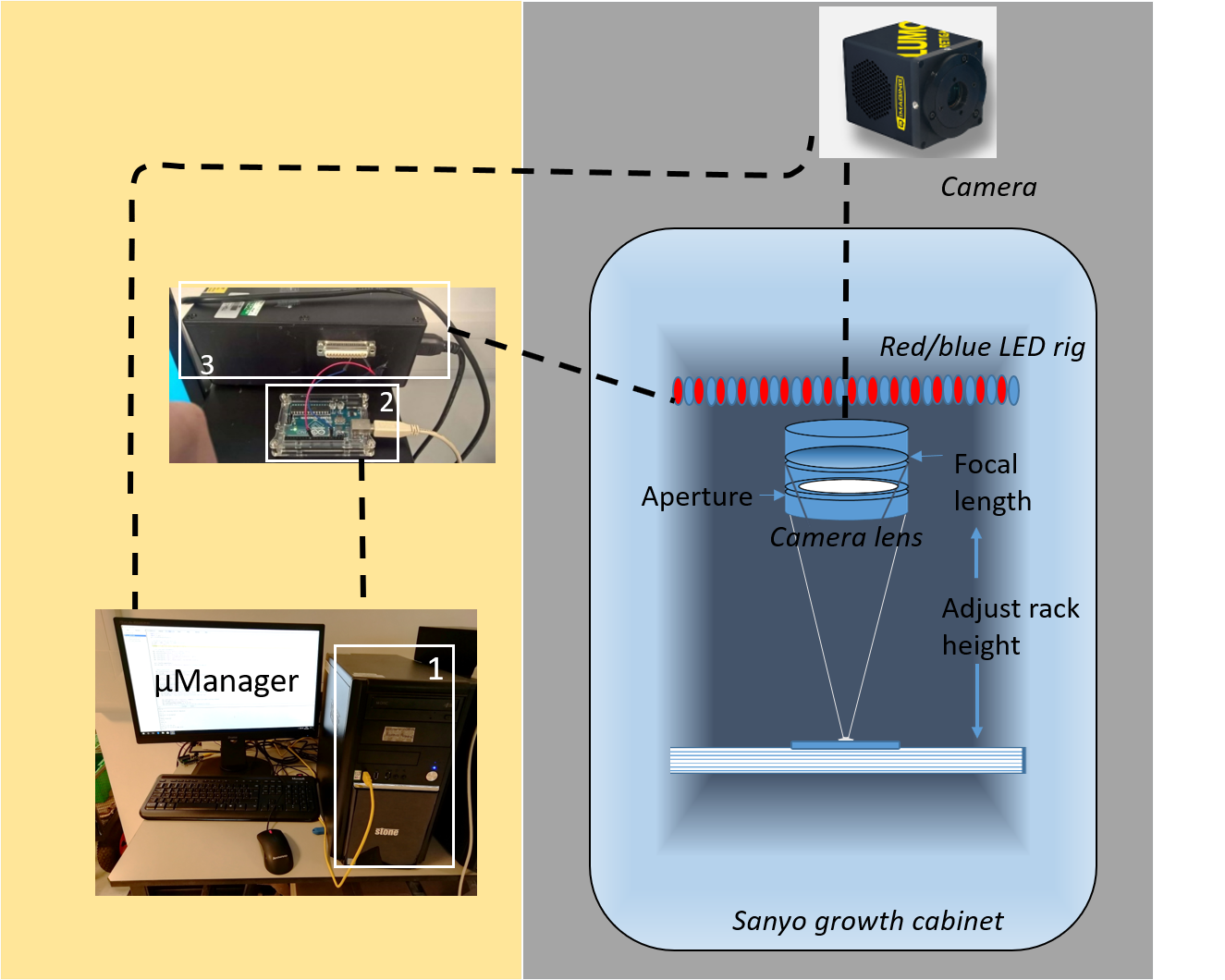

Additional Experiments

Independent experiments to those shown in the main figures. Any method differences have been highlighted.

**S11. Additional experiments showing rhythms in wheat under D:D or L:L scripts:**

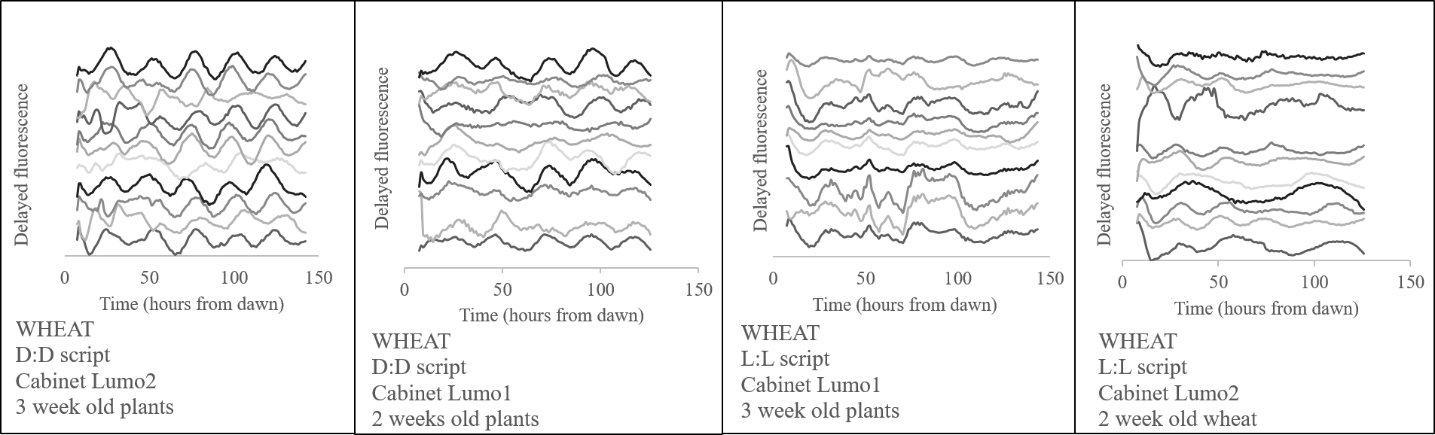
Delayed fluorescence between D:D v L:L scripts in wheat. 2 experiments- 2 week old and 3 week old wheat plants with two Lumo cameras with reciprocal tests of LL or DD script in each cabinet. Data is cubic de-trended, normalized to the mean intensity and then spread to show rhythmicity of each sample.

**S12. Additional experiments showing rhythms in *Brassica oleracia* under D:D or L:L scripts:**

Delayed fluorescence between D:D v L:L scripts in 6 different *Brassica oleracia* species. 2 experiments- 2 week old and 3 week old plants with two cameras with a L:L script used for the 2 week old experiments and the D:D script used for the 3 week old experiments. Data is cubic de-trended, normalized to the mean intensity and then spread to show rhythmicity of each sample.

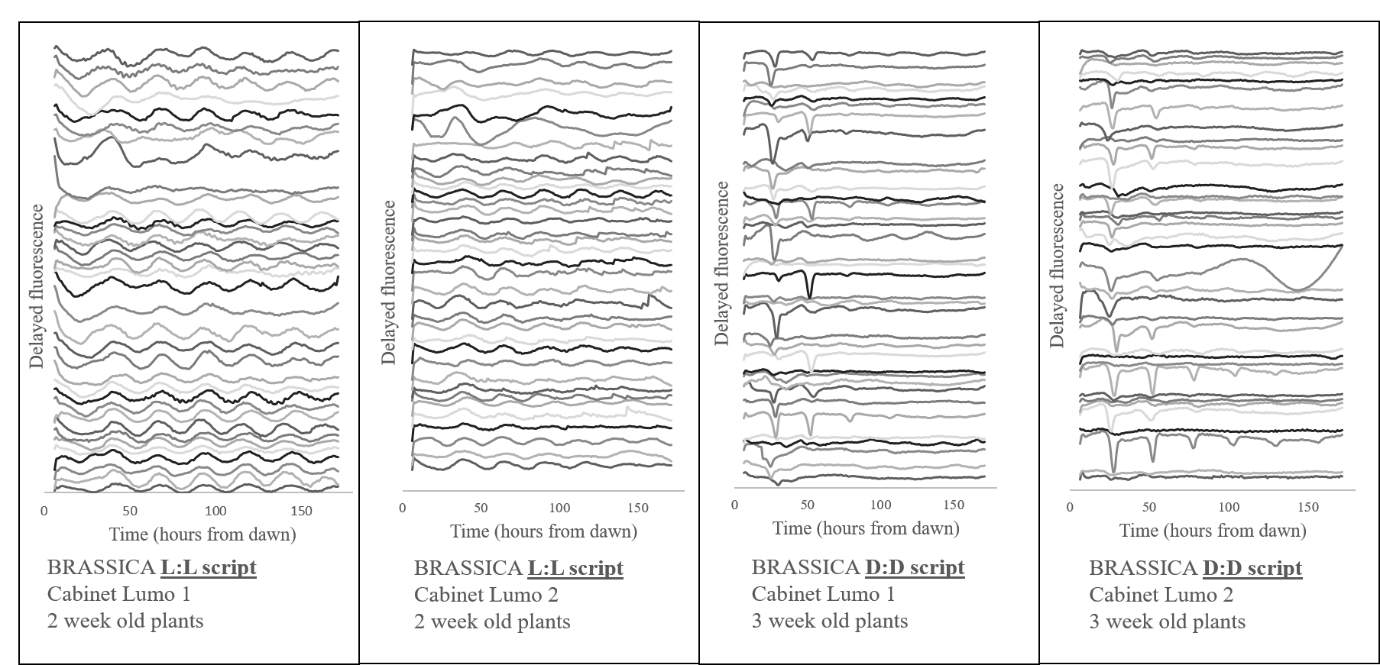

**S13. Additional experiments showing rhythms in Wheat under 17°C or 22°C:**

Delayed fluorescence in wheat under 17C or 22C temperature entrainment and imaging conditions. Temperatures were tested over 2 separate experiments using two Lumo cameras with a D:D script. Plants were either 18 or 25 days old and whole leaves were used. Data is cubic de-trended, normalized to the mean intensity and then spread to show rhythmicity of each sample.

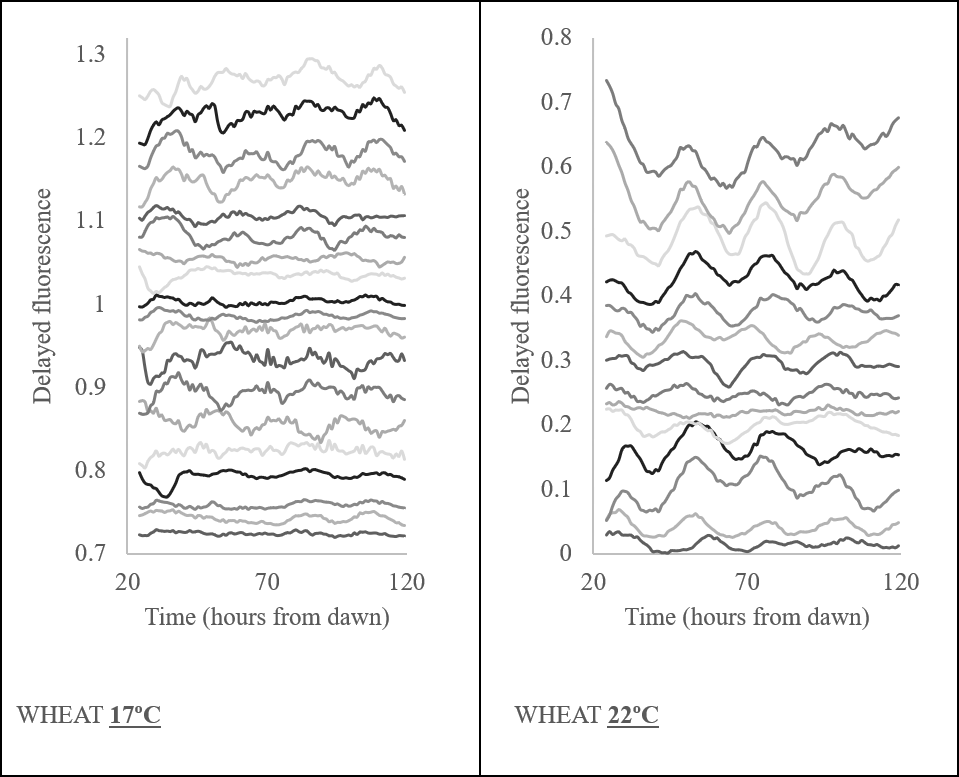

**S14: Additional experiments of the three brassica cultivars from Figure 6**

Periods detected from delayed fluorescence rhythms under L:L conditions at 22C from three independent technical replicates. Data is BAMP dtr. Cabriolet N=9/9, Chuanyou II N=7/9 and Norin N=9/9.

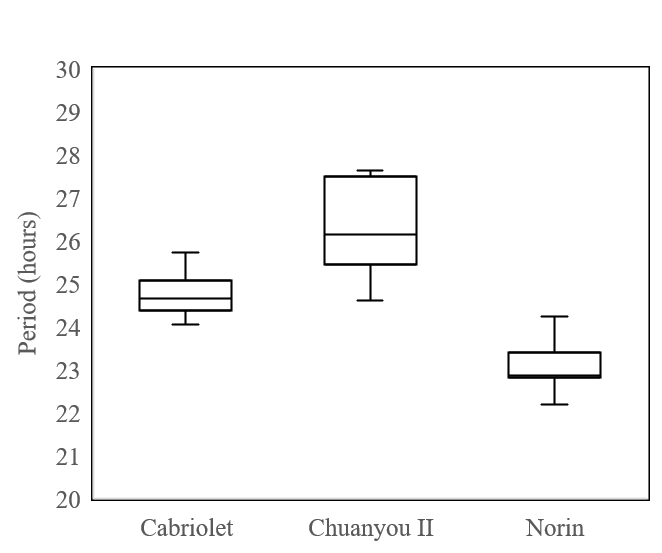
